# Investigation of the HLA locus in autopsy-confirmed progressive supranuclear palsy

**DOI:** 10.1101/2025.01.14.632901

**Authors:** Jinguo Wang, Shelley L. Forrest, Sathish Dasari, Hidetomo Tanaka, Ekaterina Rogaeva, M. Carmela Tartaglia, Susan Fox, Anthony E. Lang, Subha Kalyaanamoorthy, Gabor G. Kovacs

## Abstract

**Objectives:** Progressive supranuclear palsy (PSP) is a neurodegenerative disease showing pathological tau accumulation in subcortical neurons and glial cells. The human leukocyte antigen (*HLA*) locus on chromosome 6 is a polymorphic region with complex linkage patterns that has been implicated in several autoimmune and neurological disorders. The *HLA* locus has not been systematically examined in PSP. It is unclear whether tau and HLA can interact to induce an autoimmune disease mechanism.

**Methods:** We evaluated an autopsy confirmed PSP cohort (n=44) and compared allele/haplotype frequencies to those of the reference group of a local deceased Canadian donor pool. We performed HLA/Tau peptide binding prediction and modelling of HLA Class II and Tau Peptide interactions.

**Findings:** Odds ratio was 2.94 (95% CI 1.01 to 8.55; p=0.047) for *DQB1*\*06:01 allele, and 2.59 (95% CI 1.39 to 4.83; p=0.0025) for the narcolepsy associated haplotype (*DRB1*\*15:01*DQB1*\*06:02). One patient with 4 repeat tau PSP type pathology was a carrier of the IgLON5-associated haplotype (*DRB1*\*10:01*DQB1*\*05:01). HLA/Tau peptide binding prediction and modelling of HLA Class II/Tau Peptide interactions revealed strong binding tau peptides but not the PSP protofilament fold for alleles DQA1\**01:02DQB1\**06:02 and DQA1\**01:03DQB1\**06:01.

**Conclusion:** Our study suggests that epitopes within the tau peptide may bind to HLA alleles that are found in a subset of PSP patients supporting the notion of an autoimmune pathophysiological component. These findings have implications for subtyping and stratifying patients for therapies, including those targeting immune modulation.

## 1. Introduction

Progressive supranuclear palsy (PSP) is a neurodegenerative disease showing pathological tau accumulation in subcortical neurons and glial cells (Roemer, et al., 2022). Although the frequency of clinical PSP is lower than Parkinson’s disease (PD) in movement disorders clinics, autopsy studies suggest a much higher PSP prevalence (Driver-Dunckley, et al., 2023, Kovacs, et al., 2013). The clinical phenotypes associated with PSP brain pathology mostly belongs to the group of atypical parkinsonism or frontotemporal dementia (Hoglinger, et al., 2017). Although PSP is considered as 4-repeat (4R) tauopathy, some neuropathological features overlap with those described in the 3R and 4R tauopathy, postencephalitic parkinsonism (Jellinger, 2009) and IgLON5 autoimmune encephalitis related tauopathy (Gelpi, et al., 2024). The pathogenesis of PSP is unclear; the central role of tau is the leading hypothesis. Indeed, recent cryogenic electron microscopy (Cryo-EM) studies highlight a distinct filament fold of tau in PSP (Shi, et al., 2021). Moreover, several posttranslational modifications of tau in PSP are discussed, including tau truncation that can be interpreted as either that it follows the misfolding of tau or that truncation leads to misfolding (Rosler, et al., 2019). Importantly, genetic, positron emission tomography (PET) and bodily fluid-based studies highlight the role of the immune-mediated components in addition to the involvement of mitochondrial pathways and protein processing systems (Rosler, et al., 2019). The human leukocyte antigen (*HLA*) locus on chromosome 6 is a polymorphic region with complex linkage patterns that has been implicated in several autoimmune disorders and plays a role in the adaptive immune system (Sumitran-Holgersson, 2008). Among neurodegenerative diseases, the *HLA* locus has been mainly examined in PD and related diseases (Yu, et al., 2021, Yu, et al., 2023) or Alzheimer’s disease (AD)(Steele, et al., 2017), revealing various risk allele/haplotypes. Also, a recent study reported that AD and PD share a protective association with *HLA-DRB1*\*04 subtypes, harboring the 33H amino acid change (specifically *HLA-DRB1*\*04:04 and *HLA-DRB1*\*04:07), which have been linked to decreased neurofibrillary tangles in brains and reduced tau levels in cerebrospinal fluid (CSF). Notably, the protective *HLA-DRB1*\*04 subtypes exhibit strong binding to the aggregation-prone PHF6 sequence of tau (VQIVYK) when acetylated at K311, a key posttranslational modification (Le Guen, et al., 2023). However, none of these studies showed a robust association as seen, for example, in narcolepsy type I, a chronic sleep disorder associated with dysfunctional hypothalamic neuronal populations (Ayoub, et al., 2024), or anti-IgLON5 disease (Yogeshwar, et al., 2024, Gaig, et al., 2019). In the present study we evaluated an autopsy-confirmed PSP cohort and identified risk alleles and haplotypes, including one described in narcolepsy. Binding prediction revealed that epitopes within the tau peptide but not the misfolded PSP-tau may bind to HLA alleles that are found in a subset of PSP patients. These findings open new avenues for the interpretation of disease pathogenesis.

## 2. Methods

### 2.1. PSP sample collection

Cases showing subcortical neuronal and glial tau pathology (n=45) were selected from the University Health Network-Neurodegenerative Brain Collection (UHN-NBC, Toronto, Canada) based on the neuropathological diagnosis of PSP. Age at death, sex and neuropathologic diagnoses are provided in **Table 1**. Autopsy tissue from human brains were collected with informed consent from patients or their relatives and approval of the local institutional review board. Prior to inclusion in the study, a systematic neuropathological examination was performed following established diagnostic criteria of neurodegenerative conditions and co-pathologies (Tanaka, et al., 2024, Forrest and Kovacs, 2023).

**Table 1.** Overview of case demographics and HLA alleles. LNT: limbic predominant neuronal 4R tauopathy; PEP: postencephalitic parkinsonism (case Nr. 23 not included in the statistical analysis); PNLA: pallido-nigro-Luysian atrophy; PSP: progressive supranuclear palsy.

| Case Nr | Age | Sex | Pathology | A*1 | A*2 | B*1 | B*2 | C*1 | C*2 | DPA11 | DPA12 | DPB11 | DPB12 | DQA11 | DQA12 | DQB11 | DQB12 | DRB11 | DRB12 | DRB31 | DRB32 | DRB41 | DRB42 | DRB51 | DRB52 |
| --- | --- | --- | --- | --- | --- | --- | --- | --- | --- | --- | --- | --- | --- | --- | --- | --- | --- | --- | --- | --- | --- | --- | --- | --- | --- |
| 1 | 70 | F | PSP | 01:01 | 02:05 | 44:02 | 57:01 | 05:01 | 06:02 | 01:03 | 01:03 | 03:01 | 04:02 | 03:03 | 05:05 | 03:01 | 04:02 | 04:04 | 13:03 |  | 01:01 | 01:03 |  |  |  |
| 2 | 90 | F | PSP | 25:01 | 26:01 | 18:01 | 27:05 | 02:02 | 12:03 | 01:03 | 01:03 | 04:01 | 23:01 | 01:01 | 01:02 | 05:01 | 06:02 | 01:01 | 15:01 |  |  |  |  |  | 01:01 |
| 3 | 73 | M | PSP | 01:01 | 02:01 | 37:01 | 44:02 | 05:01 | 06:02 | 01:03 | 02:01 | 04:01 | 14:01 | 01:02 | 03:03 | 03:01 | 06:02 | 04:01 | 15:01 |  |  | 01:03 |  |  | 01:01 |
| 4 | 76 | M | PSP | 03:01 | 24:02 | 07:02 | 15:01 | 03:03 | 07:02 | 01:03 | 01:03 | 02:01 | 02:01 | 01:02 | 01:03 | 06:02 | 06:03 | 13:01 | 15:01 | 02:02 |  |  |  |  | 01:01 |
| 5 | 73 | M | PSP | 32:01 | 68:02 | 14:02 | 40:02 | 08:02 | 15:02 | 01:03 | 01:03 | 02:01 | 04:02 | 03:03 | 05:05 | 03:01 | 03:01 | 04:01 | 13:03 |  | 01:01 | 01:03 |  |  |  |
| 6 | 88 | M | PSP | 02:01 | 03:01 | 35:01 | 40:02 | 02:02 | 04:01 | 01:03 | 01:03 | 04:01 | 04:02 | 01:01 | 05:05 | 03:01 | 05:01 | 01:01 | 11:01 |  | 02:02 |  |  |  |  |
| 7 | 71 | M | PSP | 23:01 | 32:01 | 18:01 | 44:03 | 04:01 | 04:01 | 01:03 | 02:01 | 04:01 | 17:01 | 01:05 | 03:01 | 03:05 | 05:01 | 04:03 | 10:01 |  |  | 01:03 |  |  |  |
| 8 | 93 | M | PSP | 02:01 | 02:01 | 07:02 | 07:02 | 07:02 | 07:02 | 01:03 | 01:03 | 03:01 | 04:01 | 01:02 | 01:02 | 06:02 | 06:02 | 15:01 | 15:01 |  |  |  |  | 01:01 | 01:01 |
| 9 | 77 | M | PSP | 01:01 | 24:02 | 07:02 | 15:01 | 03:03 | 07:02 | 01:03 | 02:01 | 04:01 | 04:01 | 01:01 | 01:02 | 05:01 | 06:02 | 01:03 | 15:01 |  |  |  |  |  | 01:01 |
| 10 | 72 | M | PSP | 11:01 | 24:02 | 15:07 | 40:01 | 03:03 | 07:02 | 01:03 | 02:01 | 02:01 | 17:01 | 01:03 | 03:01 | 03:02 | 06:01 | 04:03 | 08:03 |  |  | 01:03 |  |  |  |
| 11 | 72 | M | PSP | 24:02 | 32:01 | 35:01 | 37:01 | 04:01 | 06:02 | 01:03 | 01:03 | 02:01 | 04:01 | 05:09 | 05:09 | 03:01 | 03:01 | 11:01 | 11:01 | 02:02 | 02:02 |  |  |  |  |
| 12 | 73 | F | PSP | 11:01 | 68:01 | 07:02 | 44:02 | 07:02 | 07:04 | 01:03 | 01:03 | 04:01 | 04:01 | 01:02 | 03:01 | 03:02 | 06:02 | 04:04 | 15:01 |  |  | 01:03 |  |  | 01:01 |
| 13 | 79 | M | PSP | 02:01 | 31:01 | 07:02 | 40:01 | 03:04 | 07:02 | 01:03 | 01:03 | 04:01 | 04:01 | 01:02 | 01:02 | 06:02 | 06:02 | 15:01 | 15:01 |  |  |  |  | 01:01 | 01:01 |
| 14 | 84 | M | LNT | 01:01 | 23:01 | 44:03 | 52:01 | 04:01 | 12:02 | 01:03 | 02:07 | 04:01 | 04:01 | 01:03 | 02:01 | 02:02 | 06:01 | 07:01 | 15:02 |  |  | 01:01 |  |  | 01:02 |
| 15 | 72 | M | PSP | 01:01 | 02:01 | 44:02 | 57:01 | 05:01 | 06:02 | 01:03 | 01:03 | 04:01 | 04:01 | 02:01 | 03:03 | 03:01 | 03:03 | 04:01 | 07:01 |  |  | 01:03 | 01:03N |  |  |
| 16 | 79 | M | PSP | 01:01 | 02:01 | 08:01 | 35:03 | 04:01 | 07:01 | 01:03 | 02:01 | 01:01 | 04:02 | 03:01 | 05:01 | 02:01 | 03:02 | 03:01 | 04:01 | 01:01 |  |  | 01:03 |  |  |
| 17 | 83 | F | PSP | 01:01 | 03:01 | 07:02 | 37:01 | 06:02 | 07:02 | 01:03 | 01:03 | 03:01 | 03:01 | 01:02 | 03:03 | 03:01 | 06:02 | 04:07 | 15:01 |  |  | 01:03 |  | 01:01 |  |
| 18 | 72 | M | PSP | 02:01 | 02:01 | 07:02 | 15:01 | 03:04 | 07:02 | 01:03 | 01:03 | 04:01 | 04:01 | 01:02 | 03:01 | 03:02 | 06:02 | 04:01 | 15:01 |  |  | 01:03 |  |  | 01:01 |
| 19 | 69 | M | PSP/PNLA | 01:01 | 03:01 | 08:01 | 14:02 | 07:01 | 08:02 | 02:01 | 02:01 | 01:01 | 05:01 | 01:02 | 05:01 | 02:01 | 06:09 | 03:01 | 13:02 | 01:01 | 03:01 |  |  |  |  |
| 20 | 87 | M | PSP | 02:01 | 26:08 | 07:02 | 51:01 | 07:02 | 15:02 | 01:03 | 02:01 | 06:01 | 10:01 | 01:01 | 01:02 | 05:01 | 06:02 | 01:01 | 15:01 |  |  |  |  |  | 01:01 |
| 21 | 69 | F | PSP/PNLA | 32:01 | 33:03 | 44:03 | 58:01 | 03:02 | 04:01 | 01:03 | 02:01 | 02:01 | 17:01 | 01:03 | 02:01 | 02:02 | 06:01 | 07:01 | 15:02 |  |  | 01:01 |  |  | 01:02 |
| 22 | 78 | F | PSP | 02:01 | 02:01 | 39:01 | 51:01 | 12:03 | 14:02 | 01:03 | 01:03 | 04:01 | 04:02 | 01:03 | 05:05 | 03:01 | 06:03 | 11:01 | 13:01 | 01:01 | 02:02 |  |  |  |  |
| 23 | 69 | M | PEP | 01:01 | 02:01 | 08:01 | 44:02 | 05:01 | 07:01 | 01:03 | 01:03 | 03:01 | 04:01 | 04:01 | 05:01 | 02:01 | 04:02 | 03:01 | 08:01 | 01:01 |  |  |  |  |  |
| 24 | 79 | M | PSP | 02:01 | 02:01 | 07:02 | 44:02 | 05:01 | 07:02 | 01:03 | 01:03 | 03:01 | 20:01 | 01:01 | 05:05 | 03:01 | 05:01 | 01:01 | 11:04 |  | 02:02 |  |  |  |  |
| 25 | 81 | M | PSP | 30:02 | 32:01 | 18:01 | 40:01 | 03:04 | 05:01 | 01:03 | 01:03 | 02:01 | 02:02 | 03:01 | 05:01 | 02:01 | 03:02 | 03:01 | 04:04 | 02:02 |  |  | 01:03 |  |  |
| 26 | 77 | F | PSP | 03:01 | 11:01 | 14:02 | 44:02 | 05:01 | 08:02 | 01:03 | 01:03 | 02:01 | 04:02 | 01:02 | 01:03 | 06:03 | 06:09 | 13:01 | 13:02 | 01:01 | 03:01 |  |  |  |  |
| 27 | 84 | F | PSP | 02:01 | 02:01 | 18:01 | 18:01 | 12:03 | 12:03 | 01:03 | 01:03 | 02:01 | 02:01 | 05:05 | 05:05 | 03:01 | 03:01 | 11:01 | 11:01 | 02:02 | 02:02 |  |  |  |  |
| 28 | 79 | F | PSP | 03:01 | 03:01 | 27:05 | 55:01 | 02:02 | 03:03 | 01:03 | 01:03 | 04:01 | 04:02 | 01:01 | 03:03 | 03:01 | 05:01 | 01:01 | 04:01 |  |  | 01:03 |  |  |  |
| 29 | 79 | M | PSP | 01:01 | 68:01 | 35:01 | 45:01 | 04:01 | 06:02 | 01:03 | 01:03 | 02:01 | 04:02 | 01:01 | 01:02 | 05:01 | 06:02 | 01:03 | 15:01 |  |  |  |  |  | 01:01 |
| 30 | 69 | M | PSP | 02:01 | 02:01 | 35:03 | 39:06 | 12:03 | 12:03 | 01:03 | 02:01 | 04:01 | 10:01 | 01:02 | 01:03 | 05:02 | 06:03 | 13:01 | 16:01 | 01:01 |  |  |  |  | 02:02 |
| 31 | 74 | F | PSP | 01:01 | 02:01 | 07:02 | 08:01 | 07:01 | 07:02 | 01:03 | 02:01 | 01:01 | 04:01 | 01:02 | 05:01 | 02:01 | 06:02 | 03:01 | 15:01 | 01:01 |  |  |  |  | 01:01 |
| 32 | 73 | F | PSP | 03:01 | 29:02 | 35:01 | 40:01 | 03:04 | 04:01 | 01:03 | 01:03 | 03:01 | 03:01 | 01:01 | 03:01 | 03:02 | 05:01 | 01:01 | 04:04 |  |  |  | 01:03 |  |  |
| 33 | 68 | M | PSP | 02:01 | 03:01 | 07:02 | 14:02 | 07:02 | 08:02 | 01:03 | 01:03 | 02:01 | 04:01 | 01:02 | 03:03 | 03:01 | 06:02 | 04:01 | 15:01 |  |  | 01:03 |  | 01:01 |  |
| 34 | 74 | F | PSP | 02:01 | 31:01 | 44:02 | 44:02 | 05:01 | 05:01 | 01:03 | 01:03 | 04:01 | 04:01 | 01:02 | 03:03 | 03:01 | 06:02 | 04:01 | 15:01 |  |  | 01:03 |  |  | 01:01 |
| 35 | 82 | F | PSP | 11:01 | 34:01 | 13:01 | 15:21 | 03:04 | 04:03 | 02:01 | 02:02 | 14:01 | 135:01 | 01:02 | 01:02 | 05:02 | 06:01 | 15:02 | 16:02 |  |  |  |  | 01:01 | 02:03 |
| 36 | 75 | F | PSP | 02:01 | 02:01 | 07:02 | 38:01 | 07:02 | 12:03 | 01:03 | 01:03 | 04:01 | 04:01 | 05:05 | 05:05 | 03:01 | 03:01 | 11:01 | 11:01 | 02:02 | 02:02 |  |  |  |  |
| 37 | 82 | F | PSP | 03:01 | 03:01 | 07:02 | 51:01 | 07:02 | 15:02 | 01:03 | 01:03 | 02:01 | 04:01 | 01:02 | 03:02 | 03:03 | 06:02 | 09:01 | 15:01 |  |  | 01:03 |  |  | 01:01 |
| 38 | 76 | F | PSP | 02:01 | 02:01 | 14:01 | 40:02 | 02:02 | 08:02 | 01:03 | 01:03 | 02:01 | 04:01 | 01:03 | 02:01 | 02:02 | 06:03 | 07:01 | 13:01 |  | 02:02 |  |  |  |  |
| 39 | 65 | F | PSP | 03:01 | 24:02 | 07:02 | 13:02 | 06:02 | 07:02 | 01:03 | 01:04 | 04:01 | 15:01 | 01:02 | 02:01 | 02:02 | 06:02 | 07:01 | 15:01 |  |  | 01:03 |  |  | 01:01 |
| 40 | 89 | M | PSP | 23:01 | 32:01 | 14:01 | 44:03 | 04:01 | 07:02 | 01:03 | 01:03 | 04:01 | 04:01 | 02:01 | 02:01 | 02:02 | 02:02 | 07:01 | 07:01 |  |  | 01:01 | 01:01 |  |  |
| 41 | 66 | F | PSP | 03:01 | 68:01 | 35:01 | 52:01 | 04:01 | 07:02 | 01:03 | 01:03 | 04:01 | 04:02 | 01:01 | 01:01 | 05:01 | 05:01 | 01:01 | 01:01 |  |  |  |  |  |  |
| 42 | 82 | F | PSP | 01:01 | 02:01 | 08:01 | 15:01 | 03:04 | 07:01 | 01:03 | 01:03 | 04:01 | 04:01 | 01:02 | 05:01 | 02:01 | 06:02 | 03:01 | 15:01 | 01:01 |  |  |  |  | 01:01 |
| 43 | 73 | F | PSP | 02:01 | 02:01 | 27:05 | 44:02 | 02:02 | 05:01 | 01:03 | 01:03 | 02:01 | 04:02 | 01:02 | 01:03 | 05:02 | 06:03 | 13:01 | 16:01 | 01:01 |  |  |  |  | 02:02 |
| 44 | 78 | M | PSP | 03:01 | 32:01 | 39:01 | 51:01 | 07:02 | 14:02 | 01:03 | 01:03 | 02:01 | 04:01 | 01:03 | 03:01 | 03:02 | 06:03 | 04:01 | 13:01 |  | 01:01 | 01:03 |  |  |  |
| 45 | 74 | M | PSP | 02:01 | 02:01 | 44:02 | 49:01 | 05:01 | 07:01 | 01:03 | 01:03 | 02:01 | 04:02 | 03:01 | 03:03 | 03:02 | 03:02 | 04:01 | 04:05 |  |  | 01:03 | 01:03 |  |  |

### 2.2. Ethics approval and consent to participate

This study was approved by the UHN Research Ethics Board (Nr. 20-5258). Autopsy tissues from human brains were collected with informed consent of patients or their relatives and approval from local institutional review boards.

### 2.3. HLA genotyping

Genomic DNA was isolated from postmortem brain tissue using a QIAGEN kit. Investigation of *HLA* locus was performed by long-range PCR (GenDx, Utrecht, The Netherlands) and followed by next-generation sequencing (NGS) using Illumina MiSeq platform. NGS data was analyzed by NGSengine V2.30 (GenDx, Utrecht, The Netherlands) with IMGT 3.52 database to obtain two-field high-resolution *HLA* typing for the *HLA-A*, *HLA-B*, *HLA-C*, *DRB1*, *DRB345*, *DQA1*, *DQB1*, *DPA1* and *DPB1* genes. For the reference HLA genotype group, samples were collected from Jan 2020 to July 2024 from deceased donors (n=1492) that donated organs to patients at UHN (Toronto, Canada) and were typed by high-resolution NGS for all HLA loci.

### 2.4. Statistical analysis

Chi^2^ and odds ratio calculations were performed with online tools (https://www.socscistatistics.com/tests and https://www.medcalc.org/calc/odds_ratio.php). False discovery rate (FDR) correction was applied using the p-values generated through chi2 statistics and the online tool https://tools.carbocation.com/FDR. For power calculations we used the G*Power tool (Ver 3.1.9.7) to run the Chi Square Statistic power analysis (Faul, et al., 2009).

### 2.5. HLA-Tau peptide binding prediction

The human Tau (2N4R) sequence was retrieved from the UniProt database (entry ID: P10636-8) to predict peptide binding to HLA class II molecules. The predictions on the peptide regions that are likely to bind different HLA alleles were performed using the NetMHCIIpan-4.3 server (https://services.healthtech.dtu.dk/services/NetMHCIIpan-4.3),(Nilsson, et al., 2023) a tool trained on binding affinity and mass spectrometry-derived eluted ligand data. To ensure comprehensive coverage the entire tau protein sequence was fragmented into overlapping 15-mer peptides offset by a single residue at the N-terminal flanking position. Binding predictions were performed for three specific alleles: DRA1\**01:01-DRB1\**15:01, DQA1\**01:02-DQB1\**06:02, and DQA1\**01:03-DQB1\**06:01. These alleles were chosen due to their higher frequency in PSP cohorts compared to controls, suggesting their potential involvement in PSP pathogenesis. Strong binding was defined as having an elution score below 1, in line with the recommendations provided by the authors of the NetMHCIIpan-4.3. To reduce computational redundancy during docking experiments, only one representative peptide was selected from overlapping 15-mer peptides sharing a binding core 9-residues or longer. This ensures that the binding core, shared among overlapping peptides, is sufficiently represented while minimizing computational redundancy.

### 2.6. Modelling of HLA Class II – Tau Peptide Interactions

To identify the potential molecular interactions that can form between the HLA-Tau peptide fragments predicted from the NetMHCIIpan-4.3 server, we performed docking studies of the top two peptides with their respective MHC class II alleles using the HADDOCK-2.4 server (Honorato, et al., 2024). Docking was conducted with default settings, ensuring reliable and reproducible results. Peptide structures were modeled using AlphaFold3 (Abramson, et al., 2024) for accurate structural representation. The coordinates for the DRA1*01:01-DRB1*\**15:01 allele were derived from the Cryo-EM structure (PDB ID: 8VRW),(Wang, et al., 2024) and for the DQA1*01:02-DQB1*\**06:02 allele, we utilized the crystal structure (PDB ID: 6DIG),(Jiang, et al., 2019) chosen based on high sequence identity. The missing residues in the crystal structure 6DIG were modelled using Modeler-10.4 software (Webb and Sali, 2016). For the DQA1*01:03-DQB1*\**06:01 allele, no suitable Cryo-EM or crystal structure was available; therefore, we modeled the 3D structure of this allele with AlphaFold3 (Abramson, et al., 2024). We performed the docking of the PSP filament (PDB ID: 7P65)(Shi, et al., 2021) and a single chain of the tau peptide from the PSP fibril to evaluate their binding affinity towards the specified alleles. Additionally, we utilized Alphafold3 to model the PSP filament peptide sequence and conducted a docking analysis to determine its potential affinity for the alleles. This we modeled to see if the PSP peptide could bind in its native conformation, which is random coil (intrinsically disordered). This analysis aimed to evaluate whether the fibril-derived peptides could bind in the HLA binding cleft and be considered potential epitopes. These approaches provided a structural framework to evaluate peptide binding and identify key interactions at the molecular level. UCSF ChimeraX (Meng, et al., 2023) was used for the analysis and to prepare the images of the docked conformations.

## 3. Results

### 3.1. Overview of neuropathology

Based on recent neuropathology criteria 44 out 45 were compatible with PSP diagnosis (**Table 1**)(Roemer, et al., 2022). Two of the 44 PSP cases showed severe pallido-nigro-Luysian atrophy pattern with less astrocytic tau pathology in the basal ganglia and cortex.(Yokoyama, et al., 2016) One further case showed the neuropathological features of limbic predominant neuronal inclusion body 4R tauopathy (Forrest, et al., 2022). Finally, one case that did not fulfil the neuropathological criteria for PSP showed 3R+4R immunoreactive neuronal tau pathology in subcortical areas together with severe nigral tau astrogliopathy with a suspected clinical diagnosis of postencephalitic parkinsonism (Tanaka, et al., 2024). The latter case was excluded from the calculation of HLA haplotype frequencies.

### 3.2. Definition of the reference group

HLA antigen (HLA-A, HLA-B, HLA-C, DRB1, DQB1 and DPB1) frequencies from UHN deceased donor local pool (n=1492) were calculated and compared to Canadian National Donor pool (n=1708), which is used for Canadian Blood Service cPRA calculation (**online supplemental file Figure S1**)(Tinckam, et al., 2015). No statistical significance was found between the two populations (p=0.8592, paired t-test). This indicates that our local donor pool is very similar to the Canadian National deceased donor pool with respect to HLA antigen level. Since Canadian deceased donor pool only has low-intermediate HLA typing, we can use the high-resolution *HLA* typing of the local deceased donor pool to serve as the PSP study control population, which was also *HLA* typed by NGS at high-resolution.

### 3.3. HLA genotypes in the PSP cohort

*HLA-DQB1*\*06:02 allele and *HLA-DRB1*\*15:01-DQB1*06:02 haplotype was over-represented in the PSP cohort (n=44) (**Table 1 and 2**). The *DQB1*\*06:02 allele frequency in the PSP cohort was 38.64%, significantly higher than the local deceased donor *DQB1*\*06:02 allele frequency of 21.98% (p=0.009, Chi^2^ test). Odds ratio for *DQB1*\*06:02 allele was 2.23 (95% CI 1.20 to 4.14; p=0.010; z-statistic: 2.545). All PSP carriers of *DQB1*\*06:02 allele were also carriers of the *DRB1*\*15:01 allele. Therefore, the *DRB1*\*15:01-*DQB1*\*06:02 haplotype frequency in our PSP cohort was also 38.64%. This haplotype frequency is nearly twice that observed in the deceased donor population, which is 19.50% (p=0.00178). Odds ratio for *DRB1*\*15:01-*DQB1*\*06:02 haplotype was 2.59 (95% CI 1.39 to 4.83; p=0.0025; z-statistic: 3.018). All these PSP patients had *HLA*-*DRB1*15:01-DQA1*01:02-DQB1*06:02* haplotype. Furthermore, *DQB1*\*06:01 allele frequency also appears to be higher in PSP patients (9.09%) vs. deceased donors (3.28%) (p=0.038). The odds ratio for *DQB1*\*06:01 allele was 2.94 (95% CI 1.01 to 8.55; p=0.047; z-statistic: 1.985). In contrast, *DQB1*\*05:01 was found to have similar frequency between PSP cases (22.73%) and deceased donors (23.93%) (p=0.853). After FDR correction, findings for *HLA- DRB1*\*15:01-*DQB1*\*06:02, *HLA-DQB1*\*06:02, and *DQB1*\*06:01 remained significant (**Table 2**).

**Table 2.** HLA alleles and haplotypes examined in the study. Values in bold indicate significant findings. n/c, not calculated.

| Allele/Haplotype | Relevance | Frequency<br>in PSP | Frequency<br>in CO | Original P<br>value (Chi2) | Benjamini-Hochberg<br>Adjusted P<br>value (Chi2) | OR (95%COI) |
| --- | --- | --- | --- | --- | --- | --- |
| DRB1*15:01-DQB1*06:02 | Narcolepsy | 0.3864 | 0.195 | **1.78e-3** | **5.35e-3** | 2.59 (1.39-4.83) |
| DQB1*06:02 | Narcolepsy | 0.3864 | 0.2198 | **9.00e-3** | **1.36e-2** | 2.23 (1.20-4.14) |
| DQB1*06:01 | - | 0.0909 | 0.0328 | **3.75e-2** | **3.75e-2** | 2.94 (1.01-8.55) |
| DRB1*04:04 | Protective in AD, PD | 0.113 | 0.0676 | 2.36e-1 | n/c | 1.76 (0.68-4.57) |
| DRB1*04:07 | Protective in AD, PD | 0.0227 | 0.0402 | 5.59e-1 | n/c | 0.55 (0.07-4.09) |
| DRB1*07:01-DQA1*02:02-DQB1*02:02 | LGI1 disease | 0.136 | 0.229 | 1.46e-1 | n/c | 0.53 (0.22-1.26) |
| DRB1*11:01-DQA1*05:01P-DQB1*03:01P | CASPR2 disease | 0.113 | 0.0904 | 3.43e-1 | n/c | 1.28 (0.49-3.32) |
| DRB1*10:01-DQB1*05:01 | IgLON5 disease | 0.0222 | 0.0275 | 8.48e-1 | n/c | 0.82 (0.11-6.12) |
| DQB1*05:01 | - | 0.2273 | 0.2393 | 8.53e-1 | n/c | 0.93 (0.45-1.91) |

In this study, we identified one case with the *DRB1*\*10:01-*DQB1*\*05:01 haplotype. The frequency of *DRB1*\*10:01-*DQB1*\*05:01 was 2.22% in the PSP cohort vs. 2.75% in the deceased donors (p=0.848). Finally, the allele frequency of *DRB1*\*04:04 (5/44 in PSP, 101/1492 in deceased donors) or *DRB1*\*04:07 (reported to be protective in AD and PD)(Le Guen, et al., 2023) (1/44 in PSP 60/1492 in deceased donors) was not significantly different between PSP cases and deceased donors (p=0.236 and p=0.558, respectively). Following amino acid alignment of all the *HLA DRB1* alleles observed in the PSP cohort, the 33H and 13H were seen in all DRB1*04 alleles (as reported previously),(Le Guen, et al., 2023) and were neither over or underrepresented in our PSP cohort vs. deceased donors (16/44 DRB1*04 in PSP and 452/1492 in the deceased donors (p=0.388). *DRB1*\*07:01-*DQA1*\*02:02-*DQB1*\*02:02 (6/44 PSP and 342/1492 deceased donors) and *DRB1*\*11:01-*DQA1*\*05:01P-*DQB1*\*03:01P (5/44 PSP and 135/1492 deceased donors) were not significantly overrepresented in PSP (p=0.146 and 0.598, respectively). In the PSP cohort, the case with limbic predominant neuronal 4R tauopathy and cases with pallido-nigro-Luysian atrophy were carriers of other rare DQB5/DQB6 haplotypes.

Power calculations for the 3 significant Chi2 statistic result, with alpha error probability set to @ 0.05, with a total sample size of 44 and two tails analysis revealed the following powers: i) DQB1*06:02: 0.772; ii) DRB1*15:01-DQB1*06:02: 0.896; iii) DQB1*06:01: 0.995.

### 3.4. HLA-Tau peptide binding prediction and modelling of interactions

The NetMHCIIpan-4.3 server predicted no strong-binding peptides for the MHC class II allele DRA1**\****01:01-DRB1\**15:01. However, two weak-binding peptides, EIVYKSPVV and LTFRENAKA, originating from the C-terminal region of the tau protein, exhibited the lowest elution scores among the predicted peptides. For the allele DQA1**\****01:02-DQB1\**06:02, one strong-binding peptide, SAKSRLQTA, derived from the P2 region of tau, was identified along with a weak-binding peptide, DHGAEIVYK, from the C-terminal region. The allele DQA1**\****01:03-DQB1\**06:01 demonstrated strong binding to two peptides: DHGAEIVYK, derived from the C-terminal region, and PGGGKVQII, originating from the microtubule-binding region of tau. Notably, the peptide DHGAEIVYK was shared between the alleles DQA1*\*01:02- DQB1\**06:02 and DQA1**\****01:03-DQB1\**06:01, highlighting its potential as a dominant epitope. These findings suggest that allele-specific preferences for peptide binding arise from distinct regions of the tau protein, with DHGAEIVYK emerging as a candidate epitope of immunological significance.

The Z-scores of peptides corresponding to their binding affinity for each allele correlate with the elution scores. However, a notable exception is observed for the allele DRA1*01:01-DRB1*\**15:01, where the Z-scores indicate potential strong binding epitopes despite the server predicting them as weak binders. Conversely, for the DQA1*01:03-DQB1*\**06:01 allele, the Z-scores of peptides are less negative, even though these peptides are identified as strong epitopes for this allele. This discrepancy in Z-scores across different alleles could be attributed to differences in the residue composition of the binding cleft. The docked conformations of the peptides with these alleles are shown in **Figure 1**. The electrostatic potential map of the binding clefts reveals that the DRA1*01:01-DRB1*\**15:01 allele has a higher proportion of negatively charged and neutral residues. In contrast, the binding clefts of DQA1*01:02-DQB1*\**06:02 and DQA1*01:03- DQB1*\**06:01 alleles are composed of neutral and positively charged residues. The hydrogen bond interactions of these peptides with alleles are shown in **Figure 2** and in the **online supplemental file Table S1**.

**Figure 1.**
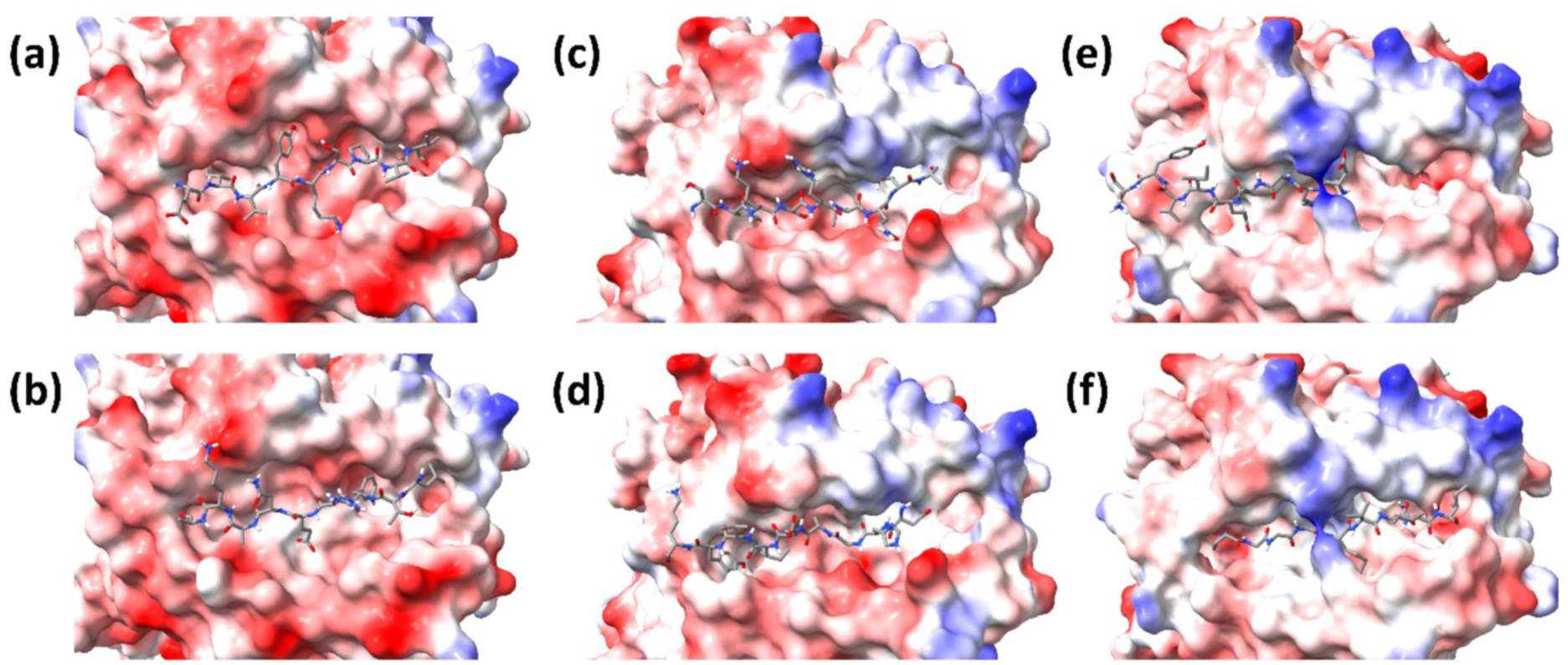
The docked conformations of the peptides with high Z-score. Please refer to Table 3 for the peptide with a high Z-score for the DRA1\**01:01-DRB1\**15:01 (a, b), DQA1\**01:02-DQB1\**06:02 (c, d) and DQA1\**01:03-DQB1\**06:01 (e, f) alleles. The alleles are shown in electrostatic map representation (Red: Negative, White: Neutral and Blue: Positive) and the peptides are shown in Licorice representation (Grey: Carbon, Red: Oxygen, Blue: Nitrogen, White: Hydrogen).

**Figure 2.**
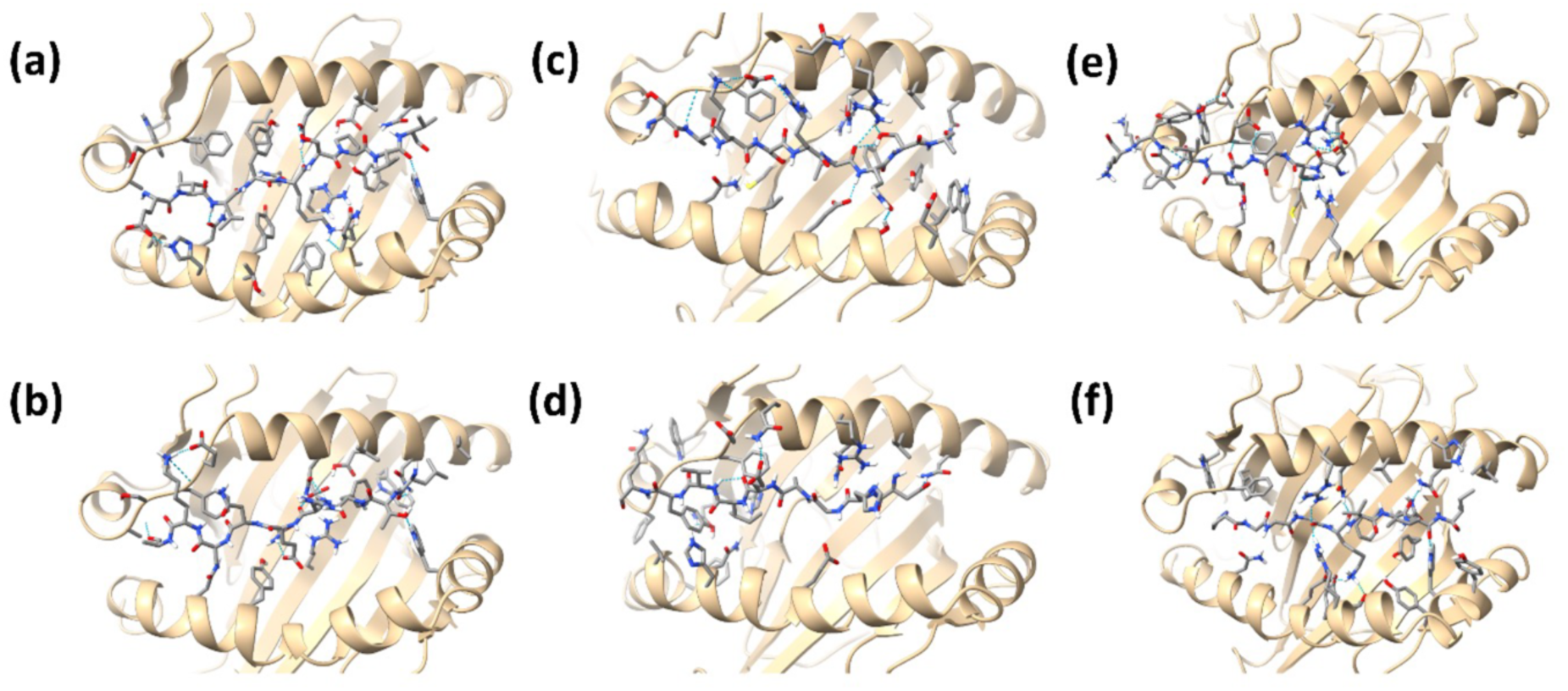
The hydrogen bond interactions formed between the peptides and DRA1\**01:01- DRB1\**15:01 (a, b), DQA1\**01:02-DQB1\**06:02 (c, d) and DQA1\**01:03-DQB1\**06:01 (e, f) alleles. Hydrogen bonds are represented with dotted cyan color.

**Table 3.** Tau peptides with low elution scores along with their Z-scores for the alleles studied.

| HLA | Full Peptide (15 AA) | Core Peptide (9 AA) | Rank EL | Binding Level | HADDOCK Z-Score |
| --- | --- | --- | --- | --- | --- |
| DRA1*01:01-DRB1*15:01 | 388-HGAEIVYKSPVVS GD-402<br>373-THKLTFR ENAKAKTD-387 | 391-EIVYKSPVV-399<br>376-LTFR ENAKA-384 | 1.32<br>1.53 | Weak<br>Weak | -81.7<br>-79.6 |
| DQA1*01:02-DQB1*06:02 | 235-SPSSAKSRLQTAPVP-249<br>384-AKTDHGAEIVYKSPV-398 | 238-SAKSRLQTA-246<br>387-DHGAEIVYK-395 | 0.79<br>1.44 | Strong<br>Weak | -70.7<br>-67.5 |
| DQA1*01:03-DQB1*06:01 | 384-AKTDHGAEIVYKSPV-398<br>267-KHQPGGGKVQIINKK-281 | 387-DHGAEIVYK-395<br>270-PGGGKVQII-278 | 0.58<br>0.67 | Strong<br>Strong | -73.6<br>-63.1 |

We also performed the docking studies of the PSP filament, single-chain peptide of the fibril and modeled PSP peptide with the alleles to evaluate their potential as epitopes. Alphafold3 predicts the PSP peptide to adopt a random coil conformation. The Z-scores representing the binding affinity of the PSP fibrils, single chain peptide of the fibril, and modeled PSP peptide with the alleles are summarized in **Table 4**. All the tested peptides exhibited very weak binding affinities towards the alleles, with the modeled PSP peptides showing the least affinity, reflected by positive Z-scores. An exception was observed for the single-chain peptide of the fibril, which demonstrated measurable binding to the DRA1\**01:01-DRB1\**15:01 and DQA1\**01:03-DQB1\**06:01 alleles. This could be attributed to the greater number of residues of the single peptide of the fibril interacting with these alleles.

**Table 4.** Z-scores of the PSP fibril, single-chain peptide of the fibril and modeled PSP peptide for the alleles studied.

| HLA | PSP | HADDOCK Z-score |
| --- | --- | --- |
| DRA1*01:01-DRB1*15:01 | Fibril | -3.6 |
|  | Single peptide of the fibril | -84.1 |
|  | Model PSP peptide | 22.8 |
| DQA1*01:02-DQB1*06:02 | Fibril | -26.5 |
|  | Single Peptide of the fibril | -16.9 |
|  | Model PSP Peptide | 27.2 |
| DQA1*01:03-DQB1*06:01 | Fibril | -10.3 |
|  | Single Peptide of the fibril | -58.4 |
|  | Modeled PSP Peptide | 6.79 |

We then evaluated docked conformations of the PSP fibril, its single-chain peptide and modeled PSP peptide with the alleles (**Figure 3**). The mode of interaction differs between the fibrils and their single-chain peptides: fibrils primarily interact through their beta-sheet structures, while the single-chain peptides engage through lateral interactions. The modeled PSP peptide interacts with the alleles through its central region. Notably, in none of these cases do the fibril, its single-chain peptide, or the modeled PSP peptide fit within the cleft of the alleles. This observation indicates a weak binding affinity, making it unlikely that they can effectively interact with the HLA alleles.

**Figure 3.**
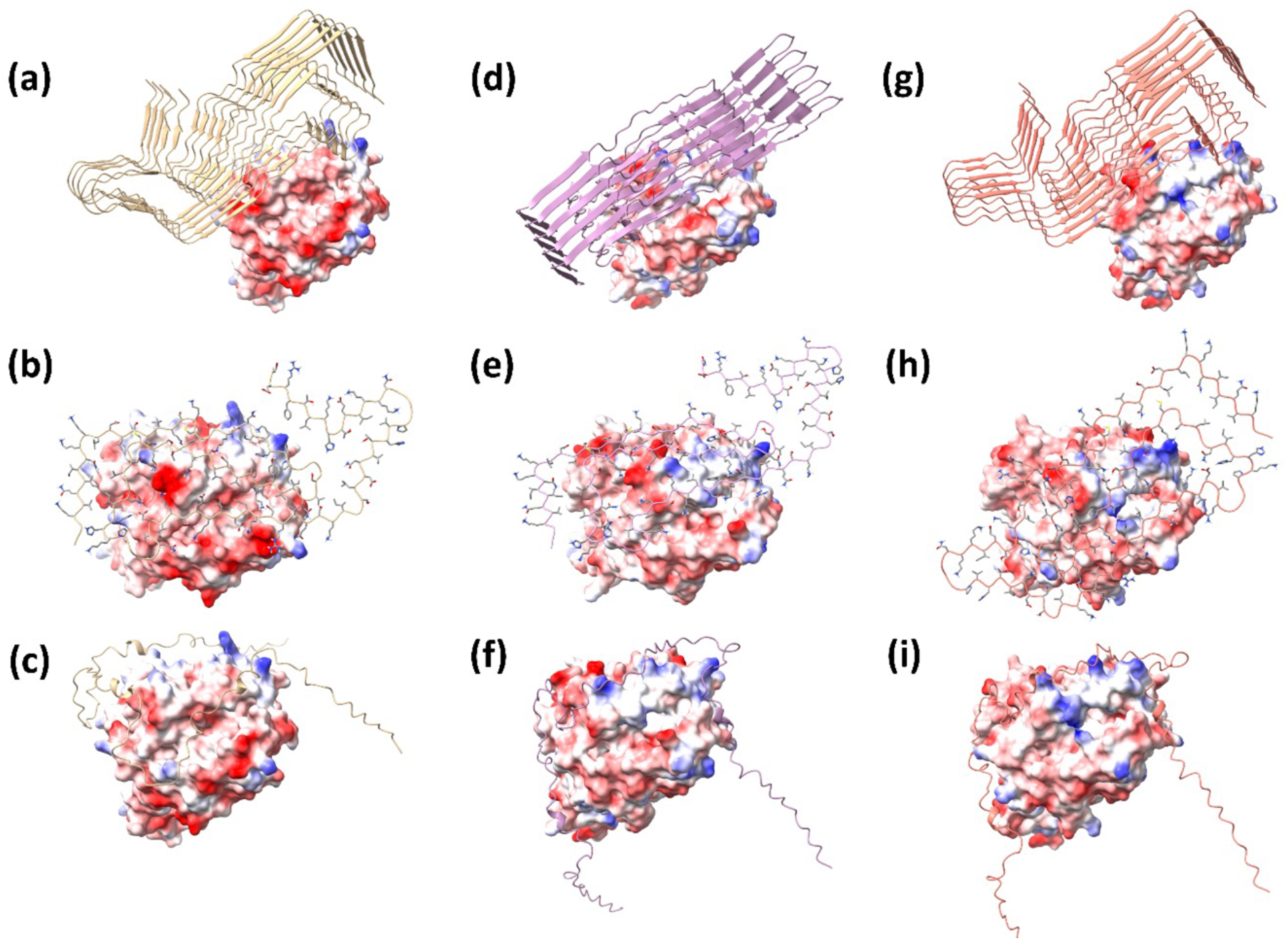
The docked conformations of the PSP filament, single chain peptide of the filament and the modeled PSP peptide for the DRA1\**01:01-DRB1\**15:01 (a, b, c), DQA1\**01:02-DQB1\**06:02 (d, e, f) and DQA1\**01:03-DQB1\**06:01 (g, h, i) alleles.

## 4. Discussion

Our study revealed that some *HLA* alleles and haplotypes are overrepresented in PSP, one of which is associated with narcolepsy. HLA-Tau peptide binding prediction and modelling of HLA Class II – Tau Peptide interactions revealed strong-binding tau peptides but not the PSP-protofilament fold for alleles DQA1\**01:02-DQB1\**06:02 and DQA1\**01:03-DQB1\**06:01. In addition, we identified one case with typical 4R PSP-type tau pathology that carried the *HLA* haplotype overrepresented in anti-IgLON5 antibody-positive autoimmune encephalitis patients (Gelpi, et al., 2024).

The *HLA* locus has not been systematically examined in PSP previously. In comparison, fine-mapping of the *HLA* locus has demonstrated a strong association of HLA-DRB1 amino acids (11V, 13H, 33H) with reduced PD risk(Yu, et al., 2021), and DR15-haplotype with increased AD risk (Steele, et al., 2017). A study on clinically defined frontotemporal dementia (i.e., that might have a wide range of neuropathological associates, including PSP) reported that 8 of the 15 identified loci mapped to the *HLA* locus on chromosome 6 (Broce, et al., 2018). Also, genome-wide association studies of PSP have indicated the presence of risk factor(s) at 6p21 (Chen, et al., 2018, Farrell, et al., 2024), including genes in the vicinity to *HLA* locus (e.g., *TNXB*)(Wang, et al., 2024). One of these studies suggested *HLA-DPB1* and *HLA-DMB* as candidate risk genes, and also found decreased RNA expression of both genes in the frontal cortex (Farrell, et al., 2024).

Our observation of overrepresentation of the haplotype associated with narcolepsy type I (i.e., *HLA-DQB1*06:02*)(Ayoub, et al., 2024) is interesting in the context that excessive daytime sleepiness, which is one clinical feature of narcolepsy (Ayoub, et al., 2024), and a different sleep disorder than that seen for example in PD (Yu, et al., 2023), can be observed in some PSP patients, although the mechanism might be different and associated with insomnia (Hattori, et al., 2003, Yasui, et al., 2006). Based on findings at the *HLA* locus, the process of autoreactivity and molecular mimicry including for viruses have been proposed (Bjornevik, et al., 2022, Luo, et al., 2018). In addition to narcolepsy, the *HLA* locus may contribute to an autoimmune origin for other brain diseases such as multiple sclerosis (*HLA-DRB1*15:01*)(Prapas and Anagnostouli, 2024), anti-IgLON5 (*HLA-DRB1*10:01-DQB1*05:01*)(Gaig, et al., 2019) and other autoimmune encephalitis syndromes associated with LGI1 (*HLA-DRB1*07:01*) and CASPR2 (*HLA- DRB1*11:01*) antibodies (Binks, et al., 2018), which were not overrepresented in our cohort. In addition to literature reports, we found one case with classical 4R PSP, and not the typical 3R+4R, tau pathology associated with the *HLA* haplotype highly associated with anti-IgLON5 disease (Gelpi, et al., 2024).

The findings at the *HLA* locus presented here differ from those found in PD and AD (Yu, et al., 2021, Steele, et al., 2017, Le Guen, et al., 2023). The recently reported AD/PD protective *HLA- DRB1*\*04 subtypes (Le Guen, et al., 2023) were not significant in our study but this result needs further evaluation in a larger cohort. Interestingly, the *DRB1*\*15:01-DQB1*06:02 haplotype that was identified with increased frequency (0.386) in our PSP cohort was also overrepresented in AD vs. controls (Steele, et al., 2017). Importantly, in the case-control study on AD (Steele, et al., 2017) the frequency of 0.189 (odds ratio = 1.08) for AD was still lower than we found for PSP in our study, and the 0.178 frequency of *DRB1*\*15:01-DQB1*06:02 haplotype in the controls of that study (Steele, et al., 2017), which is similar to our finding in deceased local donors (0.195).

The peptides with predicted low elution scores might be processed and presented by antigen-presenting cells, leading to T-cell activation as discussed also for autoimmune-mediated neurological conditions, such as narcolepsy (Ayoub, et al., 2024). Such an immune response could contribute to the breakdown of immune tolerance and the development of an autoimmune response, potentially contributing, together with other factors, to the pathology of PSP. Interestingly, the same haplotype is more frequent in AD (Steele, et al., 2017), allowing us to suggest that tau peptides may contribute to various tau-related disease when this HLA allele is present. Based on our findings we propose the concept that an autoimmune response is initiated through the presence of the rare HLA allele and the tau sequence before its aggregation process. In other words, in some cases, mis-folded PSP-tau may not be the initiator of the autoimmune response but may be a consequential downstream event. Experimental validation of these predicted epitopes is necessary to confirm their immunogenicity and clarify their role in PSP.

A further aspect is whether truncation of tau is involved in this process. Although some studies suggest that bulk truncation likely happens after assembly, however, truncation cannot be excluded in early aggregation, toxicity, or both (Goedert, et al., 2017). While many studies focused on AD, differential cleavage has been shown in PSP (Arai, et al., 2004). Truncation of tau combined with SUMO1 modification may result in a more aggregation-prone protein (Takamura, et al., 2022). Since some of the predicted peptides in our study are close to the PSP fibril-forming region, it is tempting to hypothesize that truncation might not only drive aggregation but also generate fragments with high immunogenic potential. Our concept would support that this happens before aggregation, however, for the sake of completeness it should be mentioned that some studies suggest that tau truncation may be a consequence of tau aggregation when cells attempt to eliminate protein aggregates (Wang, et al., 2010, Wang and Mandelkow, 2012), raising the possibility that the tau peptide can generate an immunogenic response without truncation.

### 4.1. Limitations

Limitations of our study are selection bias and small sample size. Given the rarity of PSP, modest sample size is an unavoidable limitation, and the selection bias is an inevitable feature of brain bank studies given the design within specialized academic centers of referral. These limitations should be considered when interpreting the results that need further confirmatory studies. However, the number of control samples we used is higher than in other studies of neurological diseases. Moreover, both our PSP and control cohorts were *HLA* typed by NGS at 4 digits (2 field) high-resolution, rather than using an imputation approach as in previous studies (Yu, et al., 2021, Steele, et al., 2017, Le Guen, et al., 2023). Since the accuracy of *HLA* genotype imputation often depends on self-reported ancestry and selected imputation algorithm,(Matern, et al., 2024) our NGS approach was more informative for our cohort. Since the *DQB1*\*06:02 allele was strongly linked to *DRB1*\*15:01 allele in our PSP cohort, we cannot further determine which of them functionally contributes to PSP. Finally, for some PSP cases we do not have information on ethnicity, which is important for predisposition to PSP (Couto, et al., 2024). Therefore, the ethnic aspect of the frequency of *HLA* alleles/haplotypes should be explored in future PSP studies.

## 5. Conclusion

Our study identifies the *DRB1*\*15:01-*DQB1*\*06:02 haplotype and *DQB1*06:01* and *DQB1*\*06:02 alleles to be strongly associated with PSP and identified potential epitopes within the tau peptide that may bind to alleles overexpressed in PSP patients raising the possibility that a subset of PSP cases might have an autoimmune pathophysiological component. This finding has important possible implications for subtyping and stratifying patients for therapies, including immune modulatory agents.

## Author Contribution

J.W., S.L.F, S.D, A.E.L, S.K., and GGK were involved in conceptualisation and methodology of the study. S.L.F., H.T., E.R., M.C.T., S.F., A.E.L, and G.G. K were part of the multicentre clinical and neuropathology team and were involved in the investigation. J.W., was performing HLA analysis and E.R., and G.G.K. were engaged in interpreting HLA the results. S.D. and S.K. performed tau peptide binding prediction and modelling of HLA Class II – Tau Peptide interactions and J.W., S.L.F, and G.G.K. were involved in interpreting results. J.W., S.L.F., S.D., E.R., S.K., G.G.K. contributed to writing of the first draft, while H.T., M.C.T., S.F. and A.E.L. contributed to the review and editing of the initial paper. All authors contributed to the writing and review of the final manuscript. J.W., S.D., E.R., S.K. and G.G.K. accessed and verified the underlying data. All authors read and approved the final version of the manuscript.

## Data sharing statement

Upon reasonable request, deidentified participant data and code used in the analyses can be shared with other researchers. Data will be made available after the ethics committee approves a study proposal and a data access agreement is signed. For further information, please contact the corresponding author.

## Acknowledgements

The authors especially acknowledge the patients and their families for their donation. GGK holds the Rossy Chair in PSP research at UHN, and ER is supported by G. Harry Sheppard Memorial Research Fund.

## Appendix A Supplementary data (1)

**Supplemental-Figure. S1.** Comparison of local deceased donor HLA frequency with Canadian National deceased donor HLA frequency at HLA antigen level.

**Supplemental-Table S1**. Key hydrogen bond interactions between the tau peptides and the alleles.

