## Supplementary material for "Investigation of the HLA locus in autopsy-confirmed progressive supranuclear palsy": online supplemental

### Supporting Information

**Supplemental-Figure. S1. Comparison of local deceased donor HLA frequency with Canadian National deceased donor HLA frequency at HLA antigen level.**

HLA antigens (HLA-A, B, C, DRB1, DQB1 and DPB1) frequency from this local donor pool (X-axis), n=1492, were calculated and compared to Canadian National Donor pool (Y –axis) which is used for Canadian Blood Service cPRA calculation.(*1*)

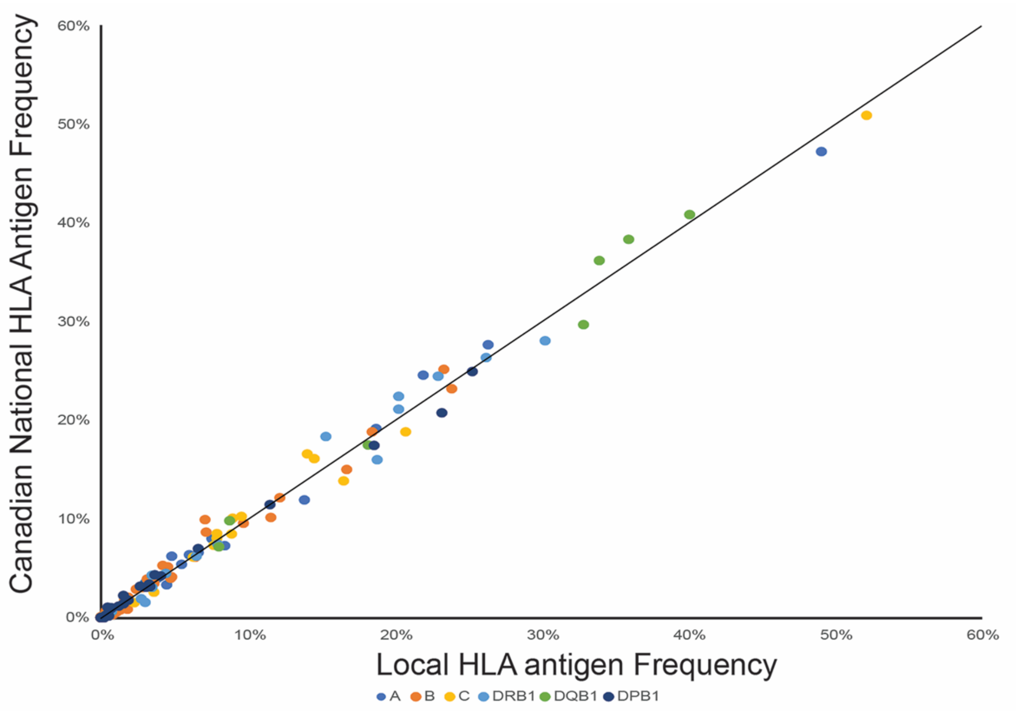

**Investigation of the HLA locus in autopsy-confirmed progressive supranuclear palsy**

Jinguo Wang, Shelley L. Forrest, Sathish Dasari, Hidetomo Tanak, Ekaterina Rogaeva, Carmela M. Tartaglia, Susan Fox, Anthony E. Lang, Subha Kalyaanamoorthy, Gabor G. Kovacs

**Supplemental-Table S1**. **Key hydrogen bond interactions between the tau peptides and the alleles.** The residue IDs for DRA1*01:01-DRB1*15:01 and DQA1*01:02-DQB1*06:02 alleles are with respect to crystal structures 8VRW and 6DIG respectively and for DQA1*01:03-DQB1*06:01 allele are with respect to the sequence excluding the signal peptide. (s: sidechain; b: backbone; left: peptide, right: allele)

| HLA | Peptide | Hydrogen bonds with alpha chain of the allele | Hydrogen bonds with beta chain of the allele |
|---|---|---|---|
| DRA1*01:01-DRB1*15:01 | EIVYKSPVV | (b) K5-N62 (s)<br>(s) S6-N62 (s) | (s) E1-H81 (s)<br>(b) V3-N82 (s)<br>(s) K5-Q70 (b)<br>(s) K5-N28 (b) |
| | LTFRENAKA | (s) K8-E55 (s)<br>(b) A9-S53 (s)<br>(s) E5-Q9 (s)<br>(b) R4-N62 (s)<br>(b) L1-N69 (s) | (s) T2-W61 (s) |
| DQA1*01:02-DQB1*06:02 | SAKSRLQTA | (b) A2-G56 (b)<br>(s) K3-F57 (b)<br>(s) K3-N58 (s)<br>(s) S4-C11 (b)<br>(s) R5-N58(s)<br>(s) T8-R64 (s)<br>(b) A9-N72 (s) | (b) Q7-E74 (s)<br>(s) Q7-T71 (s) |
| | DHGAEIVYK | (s) Y8-Q34 (S)<br>(s) E5-Q60 (b)<br>(s) D1-N72 (s) | - |
| DQA1*01:03-DQB1*06:01 | DHGAEIVYK | (b) Y8-G55 (b)<br>(s) Y8-T44 (s)<br>(s) A4-D58 (s)<br>(s) H2-C11 (b)<br>(b, s) D1-R64 (s) | (s) E5-N118 (s) |
| | PGGGKVQII | (b) Q7-N72 (s)<br>(b) K5-N65 (s)<br>(b) G4-R64 (s) | (b) I8-W97 (s)<br>(s) K5-T107 (s)<br>(s) K5-E110 (s)<br>(b) G4-R106 (s) |
